# Decapping complex is essential for functional P-body formation and is buffered by nuclear localization

**DOI:** 10.1101/2020.09.07.285700

**Authors:** Kiril Tishinov, Anne Spang

## Abstract

mRNA decay is a key step in regulating the cellular proteome. Cytoplasmic mRNA is largely turned over in processing bodies (P-bodies). P-body units assemble to form P-body granules under stress conditions. How this assembly is regulated, however, remains still poorly understood. Here, we show that the translational repressor Scd6 and the decapping stimulator Edc3 act partially redundantly in P-body assembly by capturing the Dcp1/2 decapping complex and preventing it from becoming imported into the nucleus by the karyopherin ß Kap95. Nuclear Dcp1/2 does not drive mRNA decay and might be stored there as a ready releasable pool, indicating a dynamic equilibrium between cytoplasmic and nuclear Dcp1/2. Cytoplasmic Dcp1/2 is linked to Dhh1 via Edc3 and Scd6. Functional P-bodies are present at the endoplasmic reticulum where Dcp2 potentially acts to increase the local concentration of Dhh1 through interaction with Scd6 and Edc3 to drive phase separation and hence P-body formation.

## Introduction

Translational attenuation is among the first lines of defense when a cell encounters stress. Ribosomes will release mRNA and most of the mRNA is captured into processing bodies (P-bodies), which form very quickly, within five minutes after stress encounter. The fast formation of P-bodies might be explained by the notion that the release of mRNAs and their capture into PBs is coordinated in two ways. First, a subset of PB components such as the 5’ exonuclease, Xrn1, is associated with polysomes and second, regulators of translation such as Scp160 and Bfr1 negatively regulate PB formation (Weidner *et al.*, 2014; Tesina *et al.*, 2019). Moreover, PBs contain the translational repressor Scd6, which can sequester eIF4G (Nissan *et al.*, 2010; Rajyaguru, She and Parker, 2012). P-bodies were initially thought to represent an mRNA decay compartment (Sheth and Parker, 2003). This decay requires removal of the 5’ 7-mG cap by the decapping complex Dcp1/2. The decapping activity is stimulated by DEAD box helicase Dhh1/DDX6 and the RNA binding protein Pat1 (Nissan *et al.*, 2010). Recent data, however, provide evidence that P-bodies do not only act as decay compartments but are also mRNA storage organelles and that the fate of an mRNA in PBs is dependent of the type of stressor (Wang *et al.*, 2018; Luo *et al.*, 2020). But not only the fate of mRNAs in PBs is stress-dependent, also the morphology of PBs can vary according to the stress. For example, under glucose starvation, usually 1-2 large P-bodies are observed, while hyperosmotic stress and defects in the secretory pathway induce numerous smaller P-bodies (Kilchert *et al.*, 2010). The different morphologies also suggest that the protein composition of the P-bodies might be dependent on the stressor. Although most of the research on P-bodies has been performed in *Saccharomyces cerevisiae*, because of the evolutionary conservation of the of the P-body components (Fig. 1A) and its functions, the results obtained in yeast are highly relevant for all metazoans.

**Figure 1.**
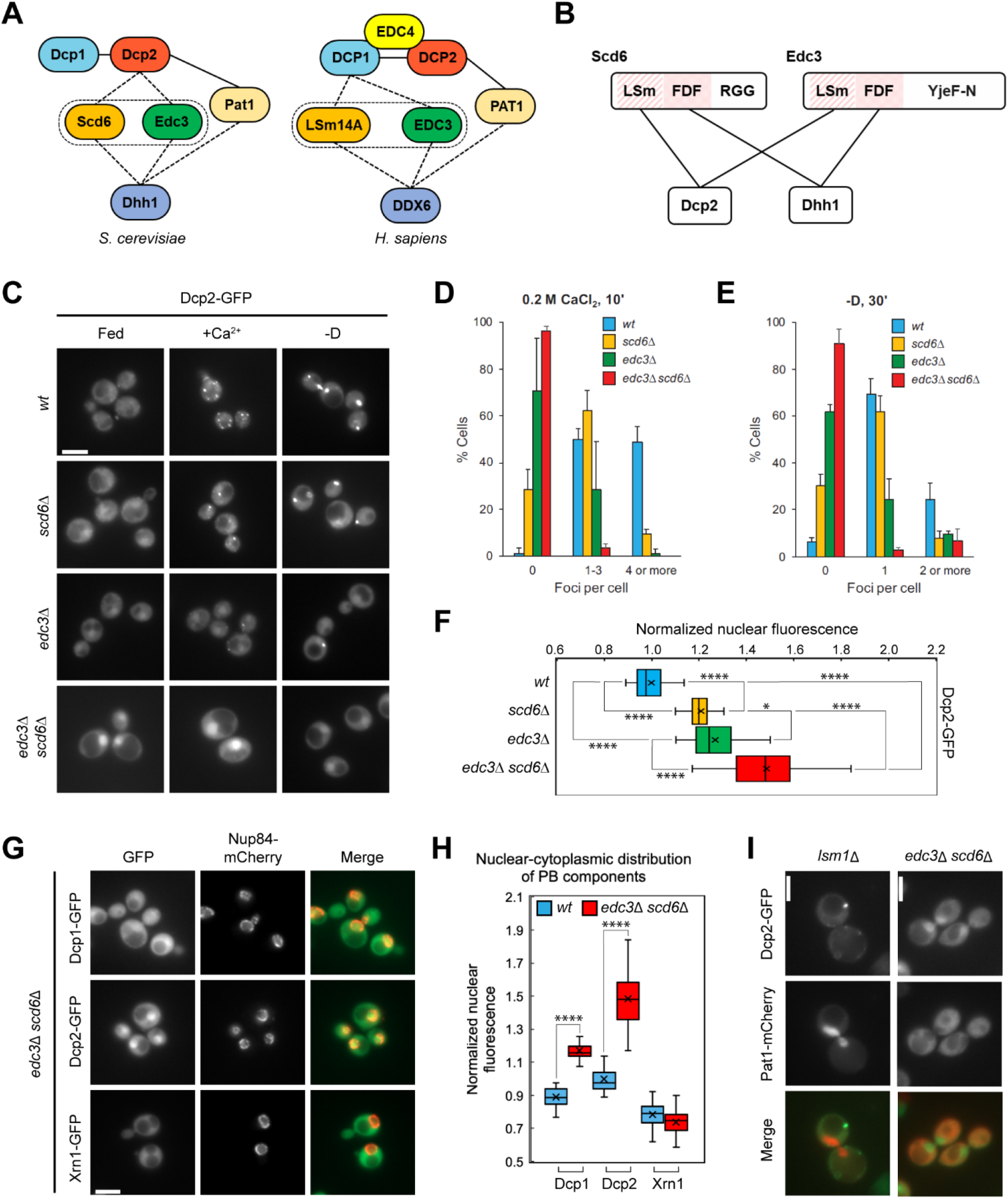
Concomitant loss of Scd6 and Edc3 blocks P-body assembly and drives nuclear accumulation of Dcp2. (A) Schematic representation of the evolutionary conserved basic PB-components in budding yeast and humans. Dashed lines indicate mutually exclusive interactions. (B) Both Scd6 and Edc3 ensure interaction between Dcp2 and Dhh1 through common structural motifs. (C) Loss of function of *SCD6* and *EDC3* leads to defects in PB formation under stress and nuclear enrichment of Dcp2. Logarithmically growing cells expressing genomically tagged Dcp2-GFP were imaged either directly or first shifted to the respective stress conditions (0.2 M CaCl_2_ for 10 min or glucose deprivation for 30 min). Scale bar 5 μm. (D) and (E) Quantification of the number of GFP foci of the data in panel C from at least 3 independent experiments. (F) Quantification of the nuclear-cytoplasmic Dcp2-GFP distribution of the data panel C from at least 3 independent experiments. The mean nuclear GFP fluorescence of a small area of the cell nucleus was normalized to the mean GFP fluorescence of small area of the cytoplasm of the same cell. (G) In *edc3*Δ *scd6*Δ the decapping complex components Dcp1 and Dcp2 are enriched in the cell nucleus while the exonuclease Xrn1 is not. Logarithmically growing *edc3*Δ *scd6*Δ cells expressing genomically tagged Dcp1-, Dcp2- or Xrn1-GFP and the nuclear marker Nup84-mCherry were imaged without additional treatment. Scale bar 5 μm. (H) Quantification of the nuclear-cytoplasmic distribution of the GFP-tagged proteins in panel H. (I) Dcp2 and Pat1 are imported into the cell nucleus through different mechanisms. Logarithmically growing cells expressing Dcp2-GFP from a genomic locus and Pat1-mCherry from a low-copy plasmid on its own promoter were imaged without additional treatment. Scale bar 5 μm.

P-bodies are membrane-less organelles. Some of the protein components contain unstructured regions or a RecA fold that are able, together with RNA to engage in liquid-liquid phase separation. Indeed, recently it was shown that Dhh1/DDX6 could drive phase separation *in vitro* and *in vivo* (Hondele *et al.*, 2019). Other proteins, such Pat1, Edc3 and Scd6 have also been shown to contribute to P-body assembly under at least some stress conditions (Decker, Teixeira and Parker, 2007; Teixeira and Parker, 2007; Kilchert *et al.*, 2010; Sachdev *et al.*, 2019). It appears as if Dhh1 and Pat1 act together, while Scd6 and Edc3 seem to have partially redundant functions in P-body formation (Coller and Parker, 2005; Decourty *et al.*, 2008; Nissan *et al.*, 2010). Thus, P-body assembly and function might be regulated through different pathways, consistent with the findings that a variety of kinases can regulate granule assembly (Yoon, Choi and Parker, 2010; Ramachandran, Shah and Herman, 2011). While under stress P-bodies are easily detected by light microscopy, they are essentially undetectable in unstressed cells. Yet, smaller P-body degradative units exist in unstressed cells as the major RNA degradation pathway in yeast is Xrn1-dependent, which acts in P-bodies (Parker, 2012). While the pathway by which Dhh1/DDX6 and Pat1 drive P-body formation is relatively well understood, information about the Scd6/Edc3-dependent pathway is still scarce.

Therefore, we decided to analyze the role of Scd6 and Edc3 in P-body assembly. We found that both proteins are required to retain the decapping complex in the cytoplasm. In an *edc3*Δ *scd6*Δ mutant, Dcp1 and Dcp2 accumulated in the nucleus through active import by Kap95. Nuclear localized Dcp1/Dcp2, however, did not drive nuclear mRNA decay. We propose that the nuclear decapping complex acts as a reservoir to regulate mRNA decay and decapping activity in the cytoplasm. We show furthermore that P-body assembly happens primarily on the endoplasmic reticulum (ER) and that in this process Edc3 and Scd6 link the decapping complex to Dhh1 (Fig.1B). Therefore, Dcp2 is also essential for P-body formation. We envisage that Dcp1/Dcp2 acts to increase the necessary critical concentration of Dhh1 at the ER to drive phase separation.

## Results

### Concomitant loss of Scd6 and Edc3 blocks P-body assembly and drives nuclear accumulation of Dcp2

Previous studies have shown that the individual deletions of *EDC3* and *SCD6* only partially affect PB formation (Decker, Teixeira and Parker, 2007; Kilchert *et al.*, 2010; Rajyaguru, She and Parker, 2012) and are dispensable for growth (Kshirsagar and Parker, 2004; Decourty *et al.*, 2008). However, since they might have partially overlapping functions, we generated an *edc3*Δ *scd6*Δ double mutant and assessed its ability to form P-bodies under stress using Dcp2-GFP as a marker (Fig. 1C). As observed previously, deletion of either *SCD6* or *EDC3* affected P-body formation under hypoosmotic stress and to a somewhat lesser extent under starvation (Fig. 1C-E) (Kilchert *et al.*, 2010). This effect was strongly exacerbated in the double mutant, consistent with the notion of a redundant function in P-body granule formation. Surprisingly, we also observed an accumulation of Dcp2-GFP in the nucleus in the single mutants, which was again strongly increased in the double mutant under normal growth conditions (Fig.1 C and F). In fact, in *edc3*Δ *scd6*Δ cells, the nuclear Dcp2 localization was maintained even under stress conditions, while this effect was less noticeable in *edc3*Δ or *scd6*Δ cells. These data suggest that there might be a correlation between Dcp2 localization and the ability of the cell to form P-body granules.

Next, we asked whether the nuclear accumulation was specific for Dcp2 or whether other P-body components would behave in a similar manner in the absence of Edc3 and Scd6. While Dcp1, which is part of the decapping complex, acted similarly to Dcp2, the exonuclease Xrn1 and the Lsm-associated protein Pat1 remained cytoplasmic (Fig. 1G-I). Therefore, loss of Edc3 and Scd6 causes the selective nuclear accumulation of the decapping complex. Nevertheless, Pat1 had been shown previously to become nuclear localized in an *lsm1*Δ mutant (Teixeira and Parker, 2007). In addition, in mammalian cells, Pat1’s interaction with both the splicing machinery in the cell nucleus and cytoplasmic P-bodies has been demonstrated (Vindry *et al.*, 2017). Yet, Dcp2 was not enriched in the nucleus under the same conditions (Fig. 1I). Our data indicate that at least two independent pathways exist to control the cytoplasmic/nuclear distribution of a subset of P-body components.

### Dcp1/Dcp2 requires active import by Kap95 for nuclear localization

It is conceivable that Dcp2 depends on Dcp1 to be localized to the nucleus in *edc3*Δ *scd6*Δ cells. To test this possibility, we generated a triple deletion *edc3*Δ *scd6*Δ *dcp1*Δ. In this strain, Dcp2 still reached the nucleus (Fig. 2A), indicating that nuclear import does not depend on the assembled decapping complex and that Dcp2 itself must contain a nuclear import signal. We analyzed the Dcp2 sequence with the NLStradamus model (Nguyen Ba *et al.*, 2009) for NLS prediction and found three potential monopartite NLSs in the amino acid regions 458-467, 562-576 and 687-708. The last region was also identified using cNLS Mapper, a web-based application for prediction of importin substrates (Kosugi *et al.*, 2009), as a part of a moderately strong bipartite NLS spanning over amino acids 673-707. The presence of putative NLSs in the Dcp2 sequence suggests the import might be mediated by the karyopherin α/β (Kap60/Kap95) complex. Both *KAP60* and *KAP95* are essential for cell viability. Therefore, we tagged *KAP95* C-terminally with an auxin-inducible degron (AID) in the *edc3*Δ *scd6*Δ background to acutely deplete Kap95 upon addition of auxin. We observed, however, that import of Dcp2-GFP into the nucleus was already impaired in the absence of auxin, indicating that the addition of the degron resulted in a hypomorphic *kap95* allele (Fig. 2B and C). Therefore, we conclude that the nuclear localization of Dcp2 is dependent on karyopherin ß.

**Figure 2.**
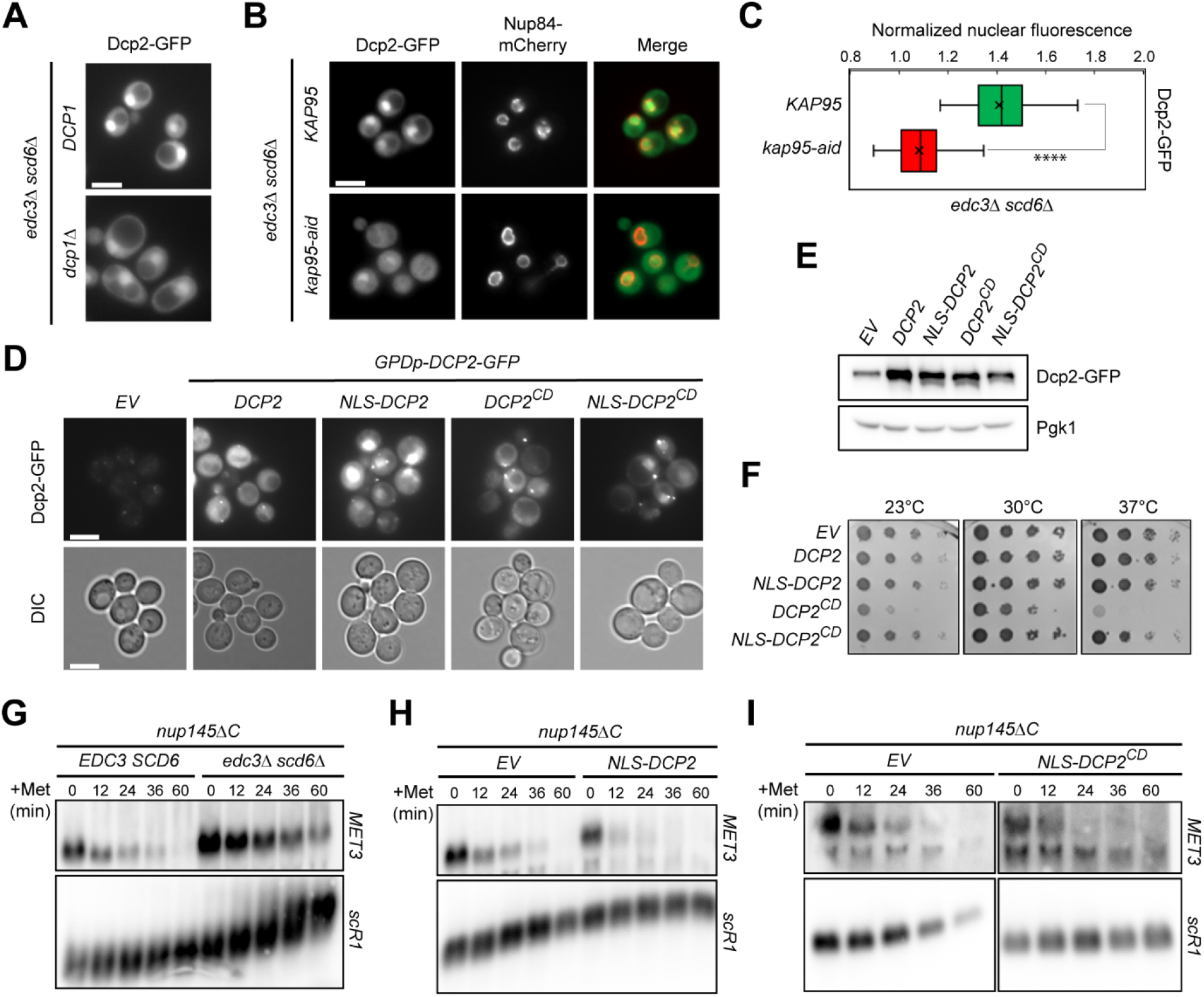
The decapping complex is stored in the nucleus as a readily releasable pool. (A) Dcp1 is not required for nuclear enrichment of Dcp2 in *edc3*Δ *scd6*Δ. Logarithmically growing cells expressing genomically tagged Dcp2-GFP were imaged without further treatment. Scale bar 5 μm. (B) Dcp2 nuclear import is dependent on Kap95. *KAP95* in *edc3*Δ *scd6*Δ cells expressing genomically tagged Dcp2-GFP and the nuclear marker Nup84-mCherry was C-terminally tagged with an auxin-inducible degron. Logarithmically growing cells from the starting strain and the *KAP95*-tagged variant were imaged without further treatment. Scale bar 5 μm. (C) Quantification of the nuclear-cytoplasmic Dcp2-GFP distribution of the data in panel B. (D) and (E) Overexpression of NLS-appended and catalytically dead (CD) variants of Dcp2. Cells expressing Dcp2-GFP from a genomic locus were transformed with a low-copy plasmid expressing different versions of Dcp2-GFP from the strong *GPD* promoter. Cells from a logarithmically grown culture were imaged by fluorescence microscopy (D, scale bar 5 μm) or GFP expression was analyzed by Western blotting (F). High levels of Dcp2 as well as Dcp2 sequestered in the cell nucleus do not affect cell fitness. Serial dilutions of the strains in panels (D) and (E) were spotted on YPD-agar, and incubated for 2 days at the indicated temperatures. (G), (H) and (I) Northern blot analysis of *MET3* mRNA nuclear degradation. *nup145*ΔC strain background was either deleted for *EDC3* and *SCD6*, or transformed with low-copy plasmids expressing the indicated Dcp2 constructs. Logarithmically growing cultures were starved for methionine to induce the expression of *MET3* and then shifted to 37°C to inhibit mRNA nuclear export. *MET3* transcription was then blocked by addition of methionine and aliquots for Northern blotting analysis were taken at the indicated time points. The *scR1* mRNA served as a loading control.

### The decapping complex is stored in the nucleus as a readily releasable pool

Next, we asked what could be the role of the decapping complex in the nucleus. Dcp1/2 could potentially decap mRNAs already in the nucleus and drive their decay there. Alternatively, the cytoplasmic concentration of active decapping complex might be tightly controlled and the nucleus would only serve as a storage space for any extra decapping complex. First, we explored a potential nuclear function of Dcp2. For this, we overexpressed GFP-tagged Dcp2 or a catalytically dead Dcp2 (Dcp2^CD^) (Van Dijk *et al.*, 2002) with or without the strong SV40 nuclear localization signal (NLS^SV40^) (Fig. 2D and E). While high levels of Dcp2^CD^ reduced cellular fitness over a range of temperatures, confining Dcp2^CD^ to the nucleus, rescued this phenotype. Likewise, a high nuclear concentration of Dcp2 was not toxic (Fig. 2F). Our data suggest that Dcp2 has essential decapping functions in the cytoplasm but not in the nucleus. Moreover, high levels of nuclear Dcp2 are well tolerated, suggesting that Dcp2 may not be active in the nucleus.

Decapped RNA is unstable in the nucleus (Kufel *et al.*, 2004). Our results above indicate that Dcp2 should not enhance RNA degradation in the nucleus. To this end, we carried out an mRNA degradation assay in a *nup145*ΔC mutant, which is deficient for mRNA nuclear export at the restrictive temperature (Kufel *et al.*, 2004). Logarithmically growing cells were first deprived of methionine at the permissive temperature (23°C) to induce the expression of the *MET3* gene and then shifted to the restrictive temperature (37°C) to block nuclear mRNA export. *MET3* expression was then shut off by addition of excess methionine to the medium, and the progress of degradation of the MET3 mRNA trapped in the nucleus was assessed by Northern blot analysis. MET3 mRNA decay was not accelerated when we trapped Dcp2 in the nucleus by either using the *edc3*Δ *scd6*Δ mutant or by overexpressing NLS^SV40^-Dcp2, or NLS^SV40^-Dcp2^CD^ as a control (Fig. 2G-I). Therefore, we conclude that Dcp2 is not actively decapping nuclear mRNAs for decay under these conditions. Taken together, our data do not support a nuclear function for Dcp2 and are consistent with the notion that the nuclear Dcp2 pool may serve as a buffer to control cytoplasmic decapping activity.

### Dcp1/Dcp2 performs essential functions on the cytoplasmic face of the ER

If the nuclear pool of Dcp2 operates as a reservoir, Dcp2’s essential function should be in the cytoplasm, in particular under stress conditions, when P-bodies are formed. Indeed, an *edc3*Δ *scd6*Δ mutant, in which Dcp2 becomes sequestered in the nucleus, showed impaired growth under stress (Fig. 3A). If this growth impairment was solely due to the Dcp2 localization, then Dcp2 overexpression (Fig. 3B and C) should rescue the growth defect. Overexpression of Dcp2 alleviated the growth phenotype of the *edc3*Δ *scd6*Δ mutant strain (Fig. 3D). These data indicate that Edc3 and Scd6 collaborate to regulate cytoplasmic Dcp2 levels and that excess Dcp2 can be stored in the nucleus until needed.

**Figure 3.**
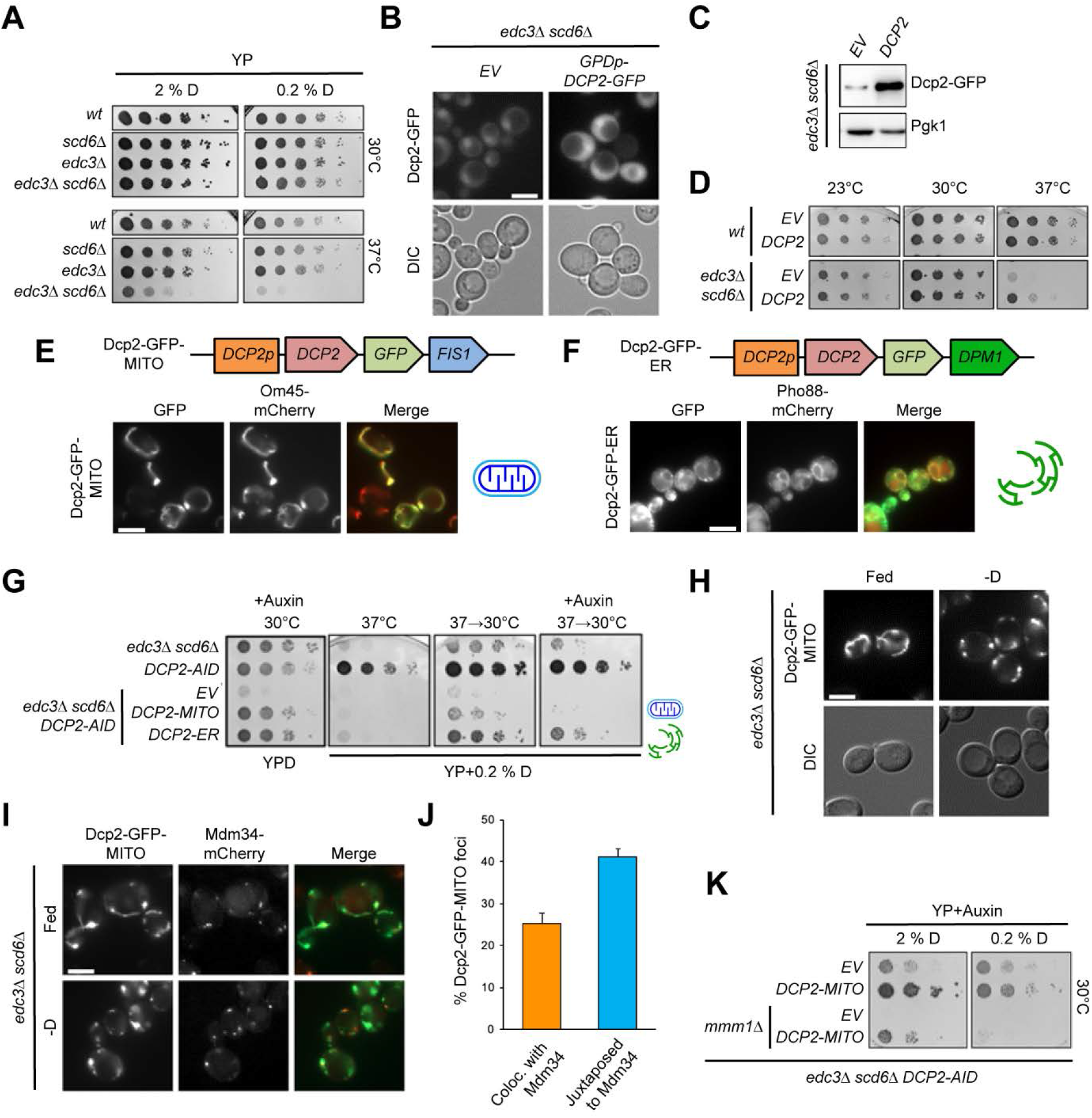
Dcp1/Dcp2 performs essential functions on the cytoplasmic face of the ER. (A) *edc3*Δ *scd6*Δ displays impaired growth under stress. Serial dilutions of the indicated strains were spotted on YP-agar containing 2 or 0.2 % glucose, and incubated at 30°C or 37°C for 2 days. (B) and (C) Dcp2 overexpression in *edc3*Δ *scd6*Δ. Dcp2-GFP was expressed on a *GPD* promoter from a low-copy plasmid on top of genomically tagged Dcp2-GFP in *edc3*Δ *scd6*Δ. Logarithmically growing cells were either imaged directly (B, scale bar 5 μm) or GFP expression was analyzed by Western blotting (C). (D) Overexpression of Dcp2 partially rescues growth of *edc3*Δ *scd6*Δ under stress. Wild-type and *edc3*Δ *scd6*Δ strains expressing Dcp2-GFP on a *GPD* promoter from a low-copy plasmid were serially diluted and spotted on YPD-agar, and incubated at the indicated temperatures for 2 days. (E) Structure and subcellular localization of the Dcp2-GFP-MITO. *edc3*Δ *scd6*Δ with genomically tagged Om45-mCherry were transformed with a low-copy plasmid expressing Dcp2-GFP-Fis1 from a *DCP2* promoter. Logarithmically growing cultures were imaged directly, scale bar 5 μm. (F) Structure and subcellular localization of the Dcp2-GFP-ER. *edc3*Δ *scd6*Δ cells with genomically tagged Pho88-mCherry were transformed with a low-copy plasmid expressing Dcp2-GFP-Dpm1 from a *DCP2* promoter. Logarithmically growing cultures were imaged directly, scale bar 5 μm. (G) Dcp2 at the ER is required for growth under stress. *edc3*Δ *scd6*Δ with *DCP2* tagged genomically with an auxin-inducible degron were transformed with low-copy plasmids expressing the indicated Dcp2-constructs from the endogenous *DCP2* promoter. Serial dilutions of the respective logarithmically growing cultures were spotted on YP-agar with 2 or 0.2 % glucose, supplemented with 0.2 M auxin as indicated. Agar plates were incubated either for 2 days at 30°C or first for 2 days at 37°C, and then for 3 days at 30°C. (H) Dcp2-MITO forms foci upon glucose starvation. *edc3*Δ *scd6*Δ strain with the endogenous *DCP2* tagged with an auxin-inducible degron was transformed with a low-copy plasmid expressing the Dcp2-MITO construct from a *DCP2* promoter. Logarithmically growing cultures were treated with 2 mM auxin for 2 h, and either imaged directly or upon 30 min glucose deprivation. Scale bar 5 μm. (I) Dcp2-MITO foci localize in close vicinity of ER-mitochondria contact sites. *edc3*Δ *scd6*Δ strain with genomically tagged Mdm34-mCherry and Dcp2-Aid was transformed with a low-copy Dcp2-GFP-MITO construct. Logarithmically growing cells were treated as for panel 3H. (J) Quantification of the number of GFP foci in panel 3-I colocalizing or juxtaposed to Mdm34 upon 30 min of glucose starvation. (K) ER-Mitochondria contact sites are essential for the functioning of Dcp2-MITO under stress. *edc3*Δ *scd6*Δ with the endogenous *DCP2* tagged with an auxin-inducible degron was deleted for *MMM1* and transformed with a low-copy plasmid expressing Dcp2-MITO from a *DCP2* promoter. Logarithmically growing cultures of the indicated strains were serially diluted and spotted on YP-agar with 2 or 0.2 % glucose, supplemented with 0.2 M auxin, and incubated at 30°C for 3 days.

The simple overexpression, however, has two drawbacks. First, it increases the entire cytoplasmic pool of Dcp2 without providing spatial information and second the overexpressed protein can still be imported and trapped in the cell nucleus. To circumvent these potential pitfalls, we locked Dcp2 in the cytosol. To this end, we anchored Dcp2-GFP on the cytosolic face of the endoplasmic reticulum (ER) and of mitochondria (mito) by linking Dcp2-GFP to Dpm1 (Dcp2^ER^) or Fis1 (Dcp2^mito^) (Fig. 3E and F). This approach allowed Dcp2 expression from its endogenous promoter and to be anchored at specific locations in the cytoplasm. We chose the ER because we have previously shown that P-bodies associate with ER membranes (Kilchert *et al.*, 2010; Weidner *et al.*, 2014; Wang *et al.*, 2018), a result that was recently confirmed in mammalian cells (Lee *et al.*, 2020). The mitochondria targeting was chosen as a control as we predicted based on our previously published results that mitochondrial Dcp2 should not be functional. The targeting approach worked as Dcp2 appended with Fis1 localized efficiently to mitochondria, and the construct carrying the Dpm1 co-localized with the ER. To ensure that the endogenous Dcp2 would not interfere with our assay, we acutely depleted endogenous Dcp2 using an auxin-inducible degron. This depletion worked efficiently as even under normal growth conditions Dcp2 depleted *edc3*Δ *scd6*Δ cells were unable to grow (Fig. 3G). Under these conditions, however, both mitochondrial and ER-localized Dcp2 rescued the growth phenotype. Under stress conditions, the ER-sequestered Dcp2 allowed much better survival when compared to Dcp2 on mitochondria (Fig. 3G). Therefore, ER-localized Dcp2 is chiefly responsible to cope with stress.

Nevertheless, we observed also some rescue by the mitochondria-localized Dcp2. To understand these results better, we first tested, whether P-bodies could be formed on mitochondria under stress conditions. Indeed, Dcp2^mito^ formed foci, resembling P-bodies under glucose starvation (Fig. 3H). These foci were also positive for two other bona fide P-body components, Dhh1 and Pat1 (Suppl. Fig. 3B and C), indicating that P-bodies can also form on mitochondria. Moreover, our data suggest that Dcp2 alone is sufficient to determine the location of P-body formation.

The ER and mitochondria are connected via contact sites to allow the exchange of lipids and ions (Elbaz and Schuldiner, 2011; Prinz, 2014). In yeast, the tethering complex ERMES stabilizes these contacts (Kornmann *et al.*, 2009). We wondered whether the P-bodies formed containing Dcp2^mito^ would be localized close to ER-mitochondria contact sites. Indeed, P-bodies were detected next to or at the same site as the ERMES component Mdm34 (Fig. 3I and J). This presence at ER-mitochondria contact sites was essential for the ability of Dcp2^mito^ to mount an appropriate stress response because destruction of the contacts by deleting the ERMES component Mmm1 abolished growth of Dcp2^mito^ expressing cells under stress (Fig. 3K). Taken together, our data are consistent with the notion that functional P-body formation under stress takes place at the ER. Moreover, our data provide evidence that Dcp2 and P-bodies can also act in trans to cope with stress at ER-mitochondrial contact sites, albeit somewhat less efficiently.

### Scd6 and Edc3 bridge the interaction of Dcp2 with Dhh1 during P-body assembly

Our data above suggest that Dcp2 localization determines where P-bodies assemble. Dcp2 can interact with both Scd6 and Edc3 (Fig. 4A), which in turn also interact with Dhh1, although their binding to Dhh1 was reported to be mutually exclusive (Fromm *et al.*, 2012). Importantly, Dhh1 has been shown to drive phase separation, which is essential during P-body formation (Mugler *et al.*, 2016; Hondele *et al.*, 2019). First, we wanted to determine which parts of Scd6 and Edc3 are required to keep Dcp2 in the cytoplasm and to promote functional stress response. To this end, we carried out a domain analysis of Scd6 and Edc3 by overexpressing their individual domains alone or in combination in the *edc3*Δ *scd6*Δ background and determined the nuclear-cytoplasmic distribution of Dcp2 and fitness at 37°C (Fig. 4B-E). As reported previously, overexpressing full-length of Scd6 impaired growth due to constitutive P-body formation, even without stress (Nissan *et al.*, 2010). Removal of the C-terminal region of Scd6 or Edc3 not only alleviated these growth defects, but restored growth to wild-type levels and reversed the P-body phenotype. Scd6 or Edc3 constructs lacking either the Dhh1 or the Dcp2 interaction site failed to restore growth and Dcp2 became enriched in the nucleus. Our data demonstrate that both Scd6 and Edc3 can bridge the interaction between Dcp2 and Dhh1, and that this connection is needed to keep Dcp2 from being transferred into the nucleus.

**Figure 4.**
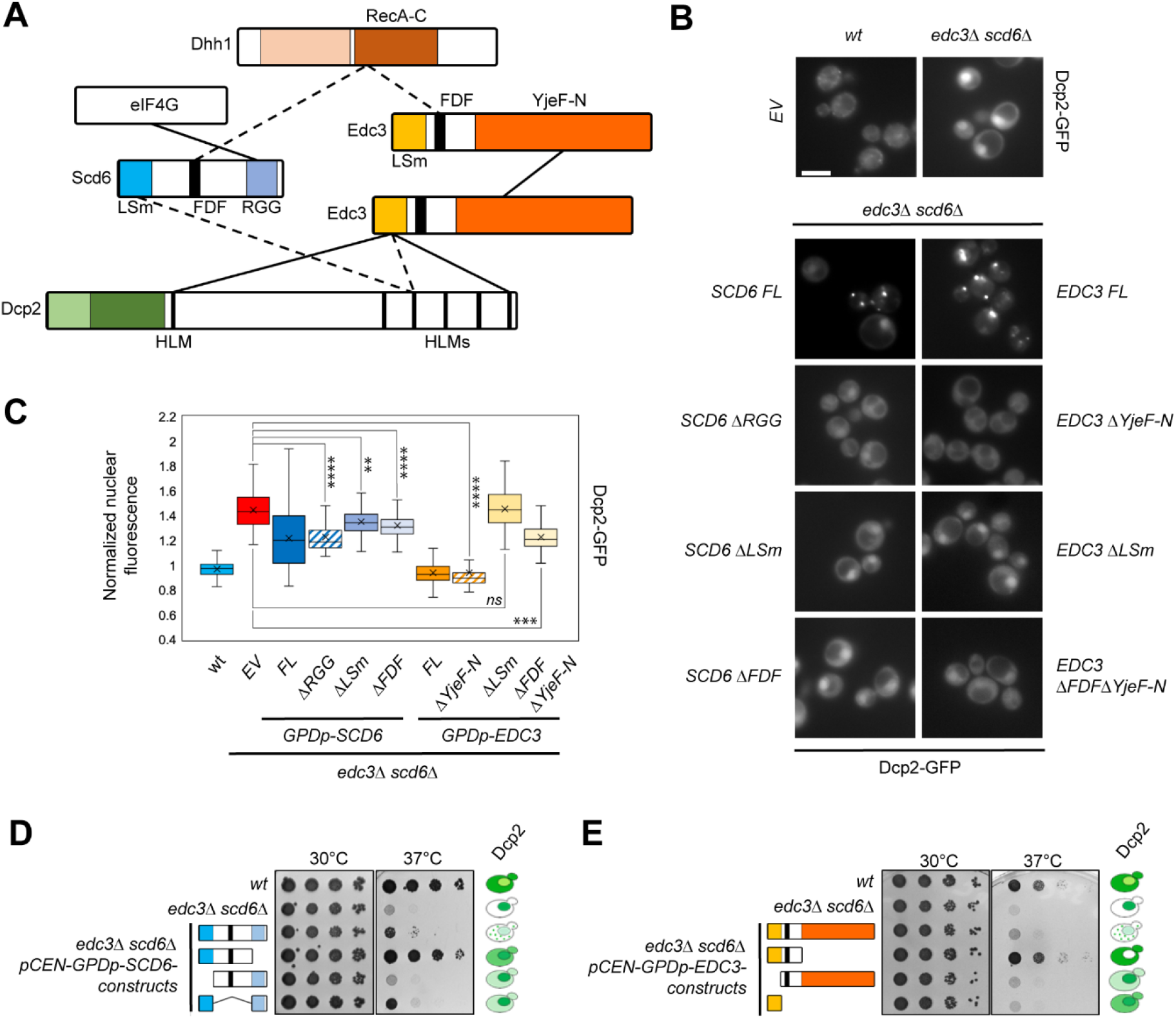
Scd6 and Edc3 bridge the interaction of Dcp2 with Dhh1 during P-body assembly. (A) Schematic representation of the domain structure and interactions of Scd6, Edc3, Dcp2 and Dhh1. Dashed lines represent mutually exclusive interactions. (B) Bridging Dcp2 to Dhh1 is required to keep Dcp2 in the cytoplasm. Wild-type or *edc3*Δ *scd6*Δ with genomically tagged Dcp2-GFP were transformed with low-copy plasmids expressing the indicated *SCD6* or *EDC3* constructs from the strong *GPD* promoter. Logarithmically growing cells were imaged without further treatment. Scale bar 5 μm. (C) Quantification of the nuclear-cytoplasmic GFP distribution in the cells on panel 4-B. D and E. Scd6 and Edc3 bridge Dcp2 to Dhh1 to cope with stress. Logarithmically growing cultures from the strains in panels B and C were serially diluted and spotted on HC-agar lacking leucine, and incubated at 30 or 37°C for 2 days. The cartoon shows the Dcp2-GFP nuclear-cytoplasmic distribution.

### Linking Dcp2 and Dhh1 drives P-body formation and functional stress response

So far, our data suggest that Dcp2 and Dhh1 must come together to drive P-body formation under stress and that this process is mediated through interaction with either Scd6 or Edc3. To further corroborate our findings, we fused the N-terminal Dcp2 binding domain of Edc3 (Edc3(1-86), Edc3^N^) to Dhh1 and expressed the construct in the *edc3*Δ *scd6*Δ Dcp2-GFP strain (Fig. 5A). Consistent with the data described above, Dcp2 is mostly nuclear in *edc3*Δ *scd6*Δ cells (Fig. 5B). This localization did not change when either Dhh1 or the Edc3^N^ were expressed separately. However, the expression of the Edc3^N^-Dhh1 fusion protein sequestered Dcp2 in the cytoplasm. More importantly, when we repeated the experiment under glucose starvation, expression of only Edc3^N^-Dhh1 was sufficient to induce P-body formation (Fig. 5C and D). These P-bodies appear to be functional as they completely rescued the growth phenotype of *edc3*Δ *scd6*Δ (Fig. 5E).

**Figure 5.**
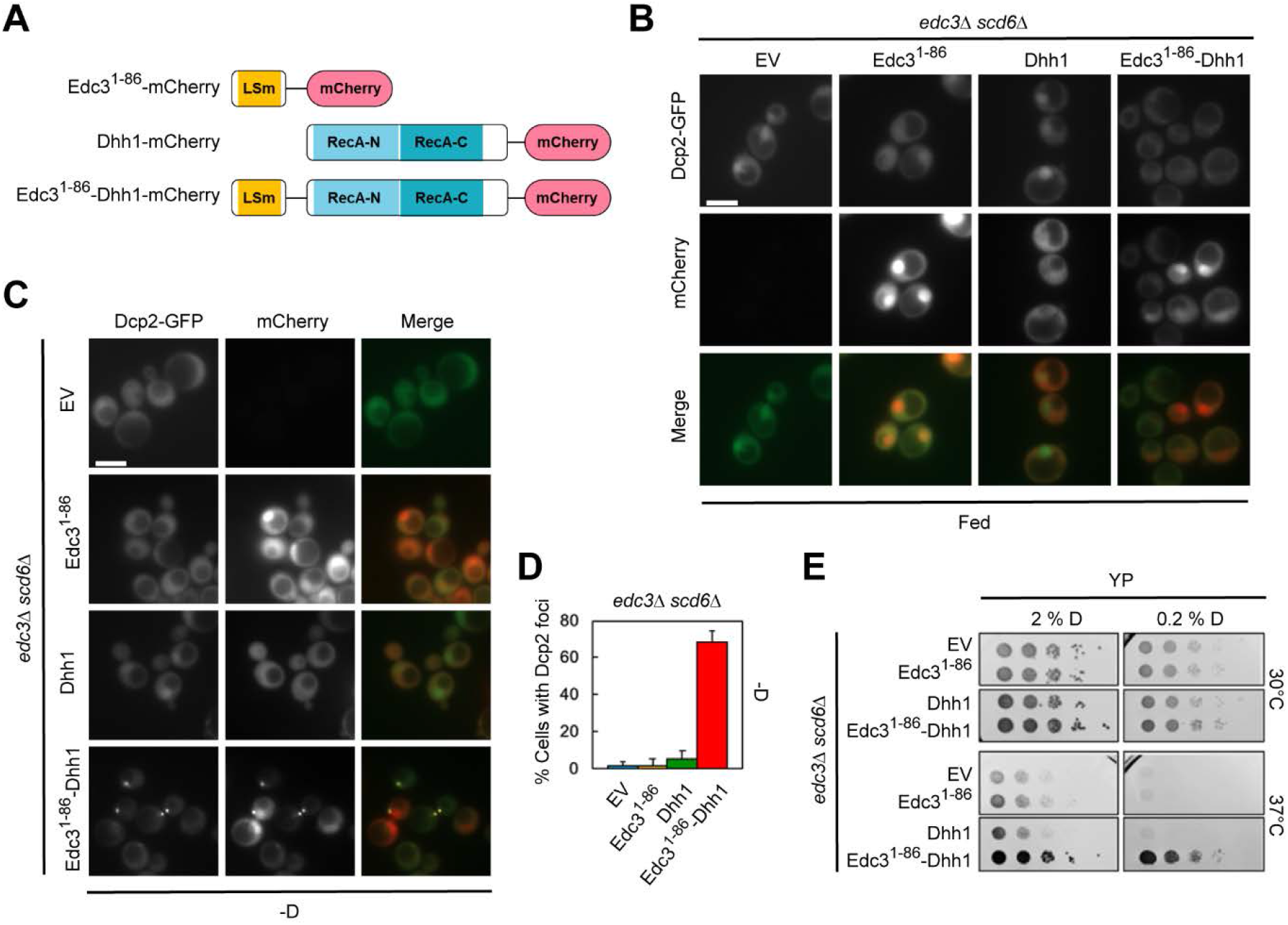
Linking Dcp2 and Dhh1 drives P-body formation and functional stress response. (A) Structure of the Edc3^1-86^-Dhh1-mCherry fusion and the control Edc3^1-86^-mCherry and Dhh1-mCherry constructs. All construct were expressed from a low-copy plasmid on the endogenous *DHH1* promoter. B-D. Linking Dhh1 to Dcp2 rescues Dcp2’s cytoplasmic localization and PB formation in *edc3*Δ *scd6*Δ. *edc3*Δ *scd6*Δ with genomically tagged Dcp2-GFP was transformed with the constructs from panel 5A. Logarithmically growing cells were imaged either directly (B) or after 30 min of glucose deprivation (C). Scale bar 5 μm. (D) Quantification of the number of cells forming GFP foci upon 30 min ‒D. (E) The Dcp2-Dhh1 interaction insures cell fitness under increased stress. Serial dilutions of the strains from panel B and C were spotted on YP-agar with 2 or 0.2 % glucose, and incubated at the indicated temperatures for 2 days.

### Dcp2 is essential for P-body assembly

Thus, our data suggest that Dcp2 is essential for P-body formation. However, it is assumed that P-body assembly is redundant and no single component is essential (Teixeira and Parker, 2007). Therefore, we decided to revisit the issue and determined P-body formation in our Dcp2-AID degron strain. Similarly, to what had been described previously, upon Dcp2 depletion, P-body formation was strongly impaired (Fig. 6A and B). Instead of 1-3 bright foci/cell, either only diffuse signal or multiple weak foci were observed, which were most conspicuous in the case of Edc3. To test whether these smaller foci might present smaller functional P-body units, we performed co-localization analyses. While Edc3 colocalized with Xrn1 and Pat1 very well in the presence of Dcp2, this level of co-localization dropped drastically in the absence of Dcp2 (Fig. 6C-E). Therefore, we conclude, that even though smaller speckles can be formed in the absence of Dcp2, they do not represent functional P-bodies, indicating that Dcp2 is essential for P-body formation. Moreover, our data are in accordance with previous data (Weidner *et al.*, 2014) that granule formation and phase separation do not necessarily correlate with P-body functionality.

**Figure 6.**
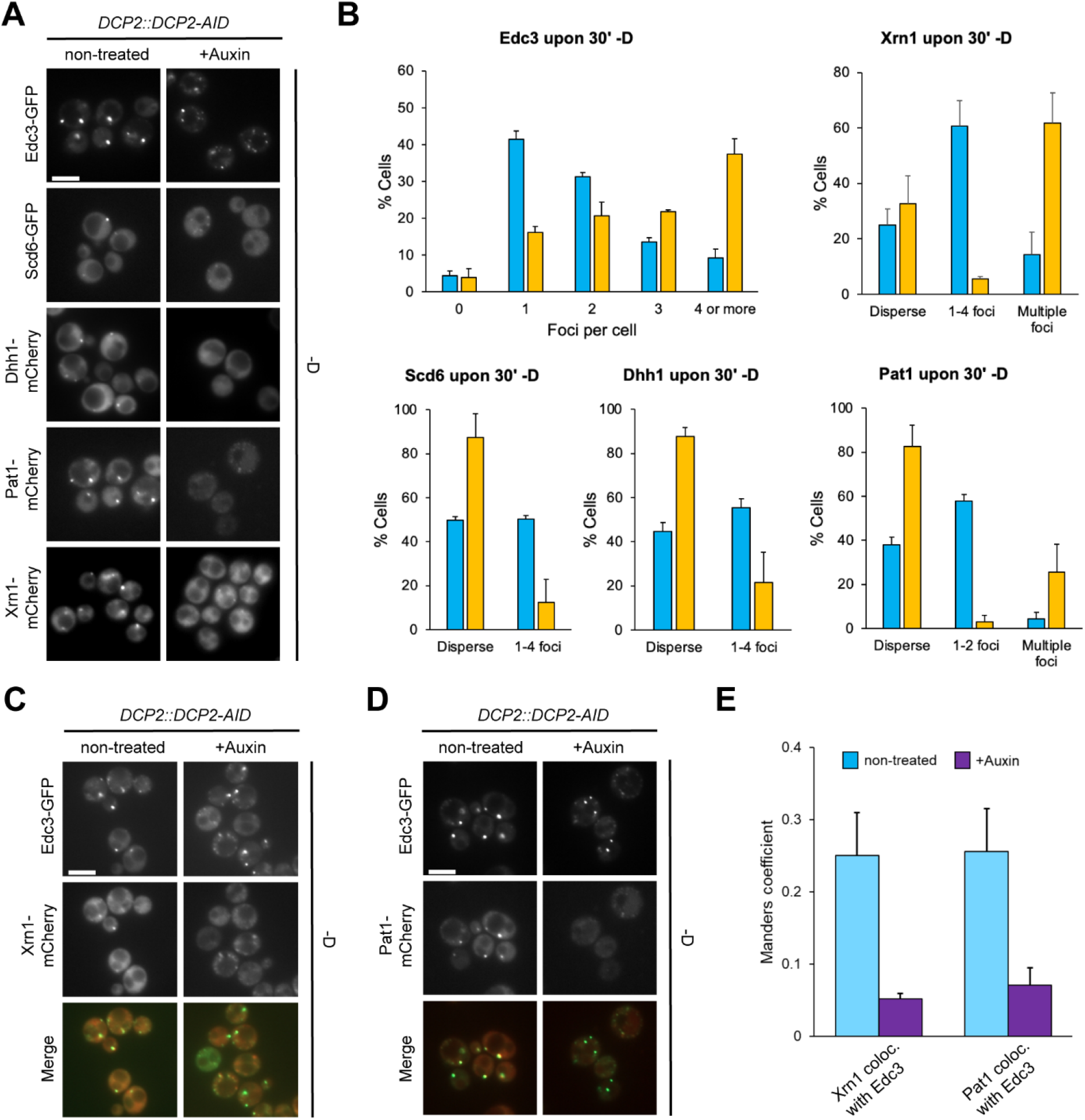
Dcp2 is essential for P-body assembly. A-B. Depletion of Dcp2 impairs PB formation. All strains were constructed from an isogenic background with *DCP2* genomically tagged with an auxin-inducible degron. The respective genes were either genomically appended with GFP or mCherry at the C-terminus (*EDC3*, *SCD6* and *XRN1*), or the respective mCherry-tagged protein was expressed from a low-copy plasmid on its endogenous promoter (Dhh1 and Pat1). Logarithmically growing cells were treated with 2 mM auxin for 2 h, then deprived from glucose for 30 min and imaged (A, scale bar 5 μm), and the number of mCherry foci was quantified (B). C-E. Dcp2 coordinates the recruitment of Xrn1 and Pat1 to Edc3 during PB formation. The strains expressing Edc3-GFP from panel 6A was genomically tagged with mCherry at the *XRN1* locus (C) or transformed with a low-copy plasmid expressing Pat1-mCherry from its endogenous promoter (D). Logarithmically growing cells were treated as in panel 6A. Scale bar 5 μm. (E) Manders coefficients for the fraction Xrn1-mCherry or Pat1-mCherry colocalizing to Edc3-GFP upon 30 min ‒D.

## Discussion

Even though P-bodies are an essential part of the cellular stress response, their assembly and cellular location are still debated and not fully understood. Recent studies highlighted the dynamics of individual P-body components during and the removal after stress (Xing *et al.*, 2020) (Lee *et al.*, 2020). The mechanism of their formation, however, is still not entirely clear. In another recent study, it was shown that the phase separation capability of the helicase Dhh1 contributed to P-body formation (Hondele *et al.*, 2019). Yet, the dogma in the field is that no individual P-body component is essential for P-body formation. This reasoning is largely based on a study in which all major P-body components were individually deleted and the formation of foci with individual P-body members was analyzed (Teixeira and Parker, 2007) with a recent follow up from the same group (Rao and Parker, 2017). The initial systematic study showed that none of the deletions suppressed completely foci formation. Here, we show that even though some P-body components form speckles in the absence of Dcp2, these appear not to be functional P-bodies as they lack other P-body components. Moreover, our data provide strong evidence that functional P-bodies are formed at the ER under stress. We envisage an assembly pathway in which Dcp2 is associated with the ER through polysomes and/or other means. We have shown previously that a phosphorylated form of Dcp2 is enriched on ER-associated polysomes under normal growth conditions (Weidner *et al.*, 2014). Moreover, immunoelectron microscopy showed P-bodies localizing in close proximity to the ER under stress (Kilchert *et al.*, 2010; Weidner *et al.*, 2014), confirmed by a recent live cell imaging approach in mammalian cells (Lee *et al.*, 2020). Another P-body component, Scd6 is associated with polysomes in the cytoplasm and on ER membranes (Weidner *et al.*, 2014), consistent with its role as a translational repressor. Upon stress, Scd6 would inhibit translation either through its direct interaction with eIF4G, or with the help of Dhh1 which likewise has translational suppressor activity (Coller and Parker, 2005; Zeidan *et al.*, 2018). However, in order to form a P-body, the local concentration of Scd6-Dhh1-mRNA complexes must be increased. This is achieved through binding to Dcp2 on the ER membrane (Fig. 7). We surmise that on the membrane, which reduces the diffusibility of the Dcp2-Scd6-Dhh1-mRNA complexes, it is much easier to gain a critical concentration to initiate Dhh1-driven phase separation. In a next step, Edc3, which like Scd6, can bind both Dcp2 and Dhh1 is recruited. Because Edc3 is a dimer, it can act as a scaffold to further enhance recruitment of Dhh1 and P-body formation. In our model, the local concentration of Dcp2 on ER polysomes would be key to drive P-body formation. In support of our model, we find that first Dcp2^ER^ can rescue a *scd6*Δ *edc3*Δ mutant, while Dcp2^mito^ cannot, unless concentrated at ER-mitochondrial contact sites. Second, directly linking Dcp2 to Dhh1 through the N-terminus of Edc3 (Edc3^N^-Dhh1) is sufficient for cells cope with stress. Moreover, an mRNA coupled to Scd6 is on the one hand translationally repressed in a Dhh1-dependent manner and on the other hand destabilized by Dcp2 (Zeidan *et al.*, 2018), supporting a temporal control in P-body assembly. In agreement with this notion, Scd6 and Edc3 interact with Dcp2 and Dhh1 in a mutually exclusive manner (Decker, Teixeira and Parker, 2007; Harigaya *et al.*, 2010; Nissan *et al.*, 2010; Fromm *et al.*, 2012; Sharif *et al.*, 2013). Finally, a recent finding did not even consider Scd6 an abundant P-body component after prolonged stress (Xing *et al.*, 2020), again supporting the idea that Scd6 plays an important role early in P-body assembly and may then be displaced by Edc3 over time. Yet, Scd6 is a bona fide P-body component because tagged Dcp2 and Scd6 are about equally efficient to purify P-bodies and to determine the RNA content upon acute stress conditions (Weidner *et al.*, 2014; Wang *et al.*, 2018).

**Figure 7.**
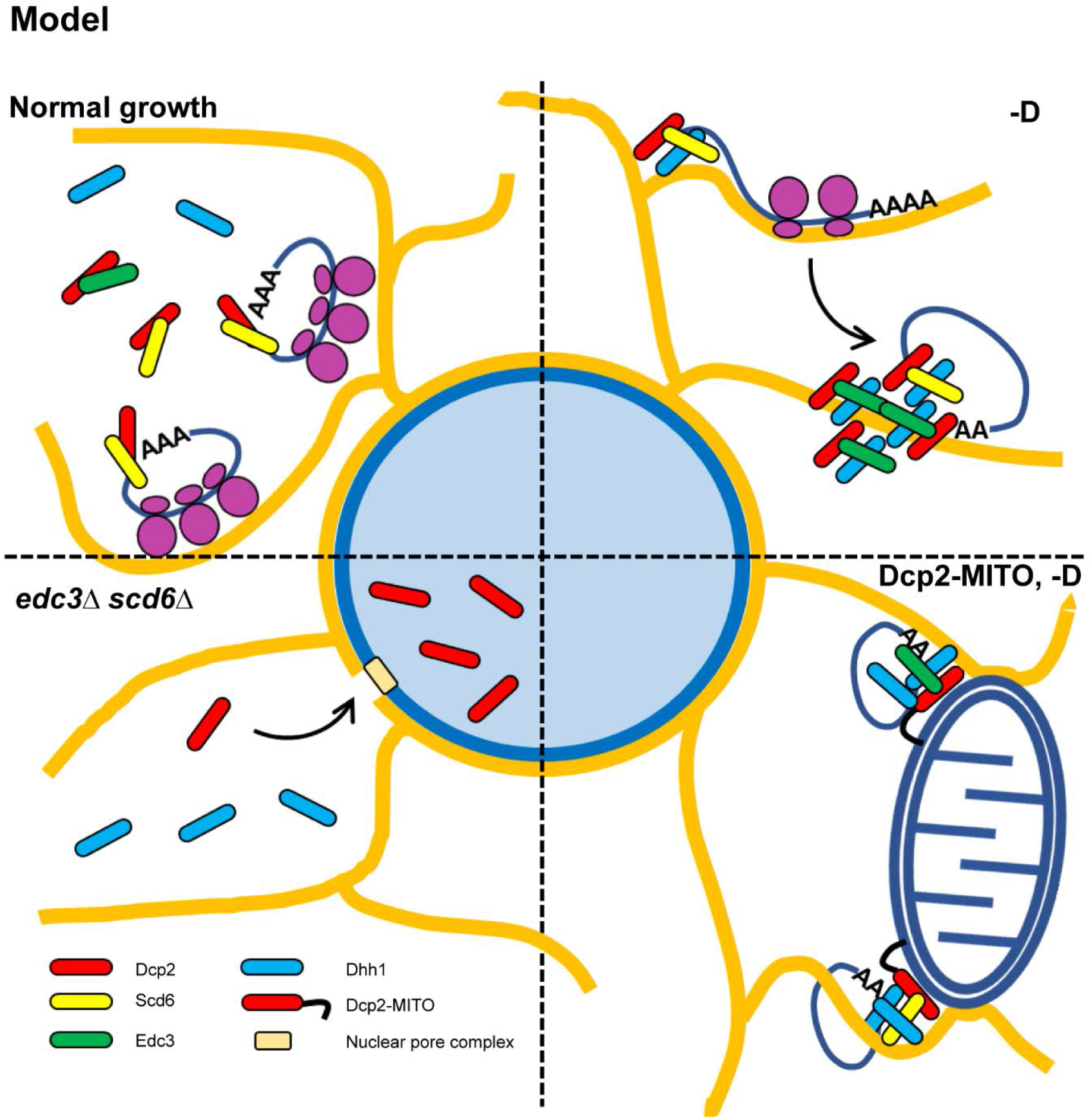
Schematic representation of our findings. For detailed information, see text.

How does our model explain why deletions of for example *PAT1* and *LSM1* also affect P-body assembly? We propose that there is an initiation phase, and this is largely the process described above in which translational repression is intimately coupled to the initiation of P-body formation. In the next step, these molecular assemblies need to grow and to be stabilized, during which process those P-body components associated with 3’ of the client mRNA are needed. In our view, in *pat1*Δ and *lsm1*Δ the initial assembly of the 5’ P-body members with the RNA at the ER is not defective, but rather stabilization of the assembly and its growth is disrupted. In this phase of P-body stabilization, the numerous partially redundant interactions among different P-body members are important as previously described (Rao and Parker, 2017). Within 5 min of stress, such as glucose deprivation, P-bodies are formed. In particular under glucose starvation, initially multiple P-bodies become visible that apparently coalesce over 30 min. Thus, P-bodies seem to mature over time, also perhaps to become more stable entities under non-adaptable stress conditions such as glucose starvation. In contrast, under adaptable stresses, this maturation process might be less critical, in particular if cells can reach a new equilibrium state within 30-45 min and start to dissociate P-bodies, as under hyperosmotic stress conditions (Kilchert *et al.*, 2010).

If Dcp2 localization, and presumably activity, are critical for P-body formation, they should be strongly controlled. We know that the activity of Dcp2 can be regulated by phosphorylation and Edc3 (EDC4 in mammals) binding (Harigaya *et al.*, 2010; Yoon, Choi and Parker, 2010; Chang *et al.*, 2014; Paquette *et al.*, 2018). Besides these on- and off switches of activity, the cell might still want to control the protein localization as a second line to control activity. Upon stress, more Dcp2 would be needed immediately to cope with the remodeling of the proteome. Acute stress demands and acute response, in which a reserve or buffering pool would be advantageous. This pool may be hidden from the cytoplasm in order to prevent premature activation and to provide a more tuned response. Dcp2 can enter the nucleus through Kap95-dependent import. The cellular/nuclear localization is determined by the presence of Scd6 and Edc3. Nuclear Dcp2 does not appear to be engaged in nuclear mRNA decay. However, it has been previously reported that nuclear Dcp2 can act as a transcriptional activator (Haimovich *et al.*, 2013). Still, the essential function of Dcp2 appears to be in the cytoplasm where overexpression of catalytically dead Dcp2 was detrimental but it had no effect when locked in the nucleus. Dcp2 is not the only P-body component that can be localized to the nucleus. The other, perhaps best documented component is Pat1 (Teixeira and Parker, 2007). Intriguingly, overexpression of Pat1 drives nuclear localization of Dhh1 (Sachdev *et al.*, 2019). It appears as if nuclear-cytoplasmic distribution of key P-body components contribute to a built-in robustness to control cytoplasmic mRNA decay.

We propose that the arrangement of the mRNA decaying factors is orchestrated around Dcp2. Dcp2 serves as a platform for organizing the different elements of the mRNA machinery on a modular principle. It associates with Scd6 and Edc3, which contact the major regulator Dhh1 responsible for liquid-liquid phase separation. An ensemble of activators (Dcp1, Edc1, Edc2, Edc3, Pat1, Lsm1-7) acts concertedly to stimulate decapping. The downstream exonuclease Xrn1 recruited to the complex by Pat1 ensures the final processing of the transcript. Liquid-liquid phase separation and condensate formation can also take place in the absence of Dcp2. The lack of the decapping platform, however, leads to loss of spatial and temporal coordination of the mRNA decay factors and failure to properly organize in P-bodies under stress.

## Material and Methods

### Yeast Methods

Strains used are listed in Supplementary Table 1. Standard genetic techniques were used throughout (Sherman, 1991). All modifications were carried out chromosomally, except where indicated. Chromosomal tagging and deletions were performed as described previously (Knop *et al.*, 1999; Goldstein and McCusker, 1999; Gueldener *et al.*, 2002; Janke *et al.*, 2004). *DCP2* and *KAP95* were genomically tagged with an auxin-inducible degron *AID*-9MYC* and *AID*-6HA* respectively using pNat-AID*-9myc and pHyg-AID*-6HA plasmid templates for generation of the C-terminal tagging cassettes (Morawska and Ulrich, 2013). *NUP145* was truncated by genomic C-terminal tagging with *3MYC* to yield the mutant *nup145*ΔC allele (nucleotides 1-1815 of the original ORF). *TRP1* marker in the genomically integrated *YIp204-ADH1p-AFB2* was disrupted using *kanMX* or *hphMX4 cassettes.*

### Plasmid construction

Shuttle vectors for expression in yeast were prepared from *pRS414-ADH* and *pRS415-GPD* backbone plasmids (Mumberg, Müller and Funk, 1995) by Gibson assembly using NEBuilder HiFi DNA Assembly Cloning Kit according to the manifacturer’s protocol (New England Biolabs). The fragments for the assembly were prepared by PCR using Q5 High Fidelity polymerase (New England Biolabs). The plasmids used along with details on their preparation are listed in Supplementary Table 2.

### Fluorescence Microscopy

Yeast cells were cultured at 30°C in YPD in case of genomically integrated modifications or in the respective HC-selection media when harboring plasmids. Cultures were diluted in HC-complete medium, re-grown to log phase, and either imaged directly or taken up in HC-medium and subjected to stresses (medium without glucose for 30 min or medium supplemented with 0.2 M CaCl_2_ for 10 min). For Dcp2 depletion by auxin-inducible degron the logarithmically growing cells before treatment with stress or imaging were supplemented with 2 mM auxin and cultured for 2 h. Fluorescence and DIC images were acquired with an ORCA-flash 4.0 camera (Hamamatsu) mounted on an Axio Imager.M2 fluorescence microscope with a 63x Plan-Apochromat objective (Carl Zeiss, Germany) and a HXP 120 C light source using ZEN 2.6 software. Image processing was performed using OMERO.insight client. For quantification of the number of foci the images from the same experiment were adjusted equally and inverted. A total of at least 300 cells from three independent experiments were quantified. Cell fluorescence measurements were carried out with ImageJ. For the nuclear-cytoplasmic GFP distribution the mean grey value of a region of interest (ROI) in the cell nucleus was normalized by the mean grey value of a ROI of the same size in the cytoplasm. A total of at least 75 cells from three independent experiments were quantified. The box and whiskers quantification graphs had the size of the box between the 25th and the 75th percentiles, and the whiskers – at the 5th and the 95th percentiles, the horizontal line marked the median and the cross indicated the mean value. The data sets were compared using a non-parametric test. *P*-values were indicated as follows: 0.1234 – ns, 0.0332 – (*), 0.0021 – (**), 0.0002 – (***), <0.0001 – (****). Colocalization was estimated using the JACoP plugin in ImageJ (Bolte and Cordelières, 2006). A total of at least 350 cells from three experiments were quantified.

### Total protein extracts and Western blot analysis

For total protein extracts 5-7 ODs of cells were spun down, resuspended in 200 μl 9 M urea, 50 mM Tris-HCl, pH 8 with freshly added 0.5 mM PMSF and beaten 2×20 s with 0.15 ml glass beads (0.25-0.5 mm) at 6.5 m/s at 4°C. 2xLaemmli buffer (200 μl) was added and the samples were denatured at 65°C for 5 min. Samples for Western blot analysis were resolved on 10 or 12.5 % SDS polyacrylamide gels and transferred on Amersham Protran Premium 0.45 μm NC membrane. Membranes were decorated with the following antibodies: rabbit anti-GFP (Torrey Pines), mouse anti-Pgk1 (Invitrogen), mouse anti-Por1 (Invitrogen), and rabbit anti-Sec61 (generous gift from Martin Spiess, Biocenter Basel), goat anti-rabbit- and anti-mouse-HRPs were from Thermo. Membranes were developed with WesternBright ECL (Advansta) at a Fusion digital imager with Evolution-capt Edge software (Vilber, France).

### Nuclear mRNA degradation assay

The nuclear degradation of the MET3 mRNA was assayed in a *nup145*ΔC mutant background. *nup145*ΔC mutants are viable at 23°C, but show a strong mRNA nuclear export defect at 37°C (Kufel *et al.*, 2004). Logarithmically growing mutant cells were first shifted to a medium lacking methionine to induce the expression of the methionine-related genes (4 h), then to 37°C for 30 min to inhibit the nuclear mRNA export and finally methionine-related genes expression was shut off by addition of excess of methionine to the medium. Aliquots were taken out, spun down and frozen in liquid nitrogen at specific times. For preparation of total RNA the cell pellets were mixed with 300 μl 50 mM sodium acetate, pH 6, 10 mM EDTA, 25 μl 20 % SDS and 300 μl phenol-chloroform-*iso*-amyl alcohol, pH 4-5. The mixtures were vortexed 30 s at top speed and incubated at 65°C for 6 min. The samples were frozen in liquid nitrogen, left to thaw for 2 min at RT and spun at 20,000xg for 10 min, 4°C. The aqueous layer was mixed with 200 μl acidic phenol-chloroform-*iso*-amyl alcohol, vortexed at top speed for 30 s and spun again. To 180 μl of the aqueous layer 20 μl 3 M sodium acetate and 600 μl ethanol was added, the mixture was chilled at −80°C for 2 h and centrifuged (20,000xg, 30 min, 4°C). The pellets were washed with 400 μl 75 % ethanol, spun down again, air-dried at RT, and dissolved in 30 μl water. An equal volume of 2X RNA Loading Dye (Thermo Fisher) was added, the samples were denatured at 65°C for 5 min and quickly chilled in an ice-water bath. For Northern blot analysis 3-5 μg total RNA were separated on a 1.2 % agarose gel containing formaldehyde. The RNA was transferred to a Hybond N+ membrane (Amersham) and hybridized to *MET3* and *scR1* digoxigenin-labelled RNA probes. The probes were prepared by *in vitro* transcription with the MegaScript T7 kit (Ambion), Digoxigenin-11-UTP (Roche) and purified DNA templates generated by PCR. One probe covering nucleobases 20-520 was used for *scR1* and two probes covering nucleobases 5-678 and 845-1446 were used in equimolar amounts for *MET3*. Membranes were decorated with sheep anti-digoxigenin Fab fragments coupled to peroxidase (Roche). The blots were developed with WesternBright ECL (Advansta) at a Fusion digital imager with Evolution-capt Edge software (Vilber, France).

### Subcellular fractionation

Logarithmically growing cells (25 ODs) were spun down, reduced in 100 mM Tris-HCl, pH 9.4, 10 mM DTT for 10 min and washed with 2×5 ml spheroplast buffer (0.7 M sorbitol, 0.7×YPD, 50 mM Tris-HCl, pH 7.5). Cells were resuspended in spheroplast buffer (50 ODs/ml), 1 mM DTT was added and the cell wall digested with Zymolyase 20T (30 μg/OD) for 20 min at RT and gentle agitation. The spheroplast suspension was then layered on a 1-ml cushion of 7.5 % Ficoll 400, 0.7 M sorbitol and spun at 1,000xg for 2 min. The spheroplast pellet was gently resuspended in 5 ml spheroplast buffer and the spheroplasts were recovered for 1 h at RT and gentle rocking. The suspensions were spun down (1,000×g, 2 min), the pellets were chilled on ice and washed with 0.8 ml ice-cold lysis buffer (0.25 M sorbitol, 20 mM HEPES, pH 6.8, 5 mM MgSO_4_, 0.5 mM PMSF). The pellets were resuspended in lysis buffer (100 ODs/ml), sonicated 2×45 s in a low-output cleaning bath at 4°C and spun at 2,000xg for 2 min, 4°C. Typically, 0.15 ml S2 was spun at 13,000xg for 10 min, 4°C. The supernatant was spun at 110,000xg for 30 min, 4°C in a TLA 100.3 rotor (Beckman). P13, P100 and S100 were recovered, denatured with Laemmli buffer at 65°C for 5 min and analyzed by Western blotting.

## Acknowledgements

We would like to thank Martin Spiess for the Sec61 antibody. We thank Rod Lim for suggesting Kap95 as responsible nuclear import factor for Dcp2. We acknowledge Maria Hondele and Ian G. Macara for critical comments on the manuscript. This work was supported by Swiss National Science Foundation (grants 310030B_163480 and 310030_185127) and the University of Basel.

## Author contribution

KT and AS conceived the study and designed the experiments. KT performed the experiments. KT and AS interpreted the data. KT wrote the first draft. AS wrote the final draft with input from KT.

